# Proteasome granular localization is regulated through mitochondrial respiration and kinase signaling

**DOI:** 10.1101/2022.01.13.476202

**Authors:** Kenrick A Waite, Jeroen Roelofs

## Abstract

In yeast, proteasomes are enriched in cell nuclei where they execute important cellular functions. Nutrient-stress can change this localization indicating proteasomes respond to the cell’s metabolic state. However, the signals that connect these processes remain poorly understood. Carbon starvation triggers a reversible translocation of proteasomes to cytosolic condensates known as proteasome storage granules (PSGs). Surprisingly, we observed strongly reduced PSG levels when cells had active cellular respiration prior to starvation. This suggests the mitochondrial activity of cells is a determining factor in the response of proteasomes to carbon starvation. Consistent with this, upon inhibition of mitochondrial function we observed proteasomes relocalize to granules. These links between proteasomes and metabolism involve specific signaling pathways, as we identified a MAP kinase cascade that is critical to the formation of proteasome granules after respiratory growth but not following glycolytic growth. Furthermore, the yeast homolog of AMP kinase, Snf1, is important for proteasome granule formation induced by mitochondrial inhibitors, while dispensable for granule formation following carbon starvation. We propose a model where mitochondrial activity promotes proteasome nuclear localization.

**Summary:** We determined a role for mitochondrial respiration in regulating proteasome granule formation and identified the cell integrity MAP kinase pathway and Snf1 kinase as regulatory factors.

## Introduction

The efficiency of many cellular processes relies on optimal levels of the proteins involved. To achieve this, there is a network of factors that coordinate protein synthesis, folding, localization, and degradation, known as the proteostasis network. Protein degradation is mainly controlled by two pathways in eukaryotes, the Ubiquitin-Proteasome System (UPS), and the autophagy-lysosome pathway (Finley *et al*., 2012; Feng *et al*., 2014; Dikic, 2017). Substrates of the UPS are labeled with one or more ubiquitins in a highly selective process (Hershko and Ciechanover, 1998; Schrader, Harstad and Matouschek, 2009; Finley *et al*., 2012). In humans more than 600 ubiquitin ligases are dedicated to the recognition and labeling of specific proteasome substrates (Li *et al*., 2008). While many ubiquitinated proteins are recognized and degraded by proteasomes, ubiquitination also serves other functions depending on the type of ubiquitination and cellular state (Fujii *et al*., 2009; Shaid *et al*., 2013; Rittinger and Ikeda, 2017).The second pathway of protein degradation is the autophagy-lysosome system. Here, macro-autophagy, hereafter referred to as autophagy, is the major pathway. Autophagy (literally self-eating) involves the targeting of proteins to the concave side of de novo formed double membrane structures. Upon expansion and closure of these structures, an autophagosome is formed that has engulfed cytosolic cargo (Kabeya *et al*., 2000; Feng *et al*., 2014; Dikic, 2017). Fusion of the outer membrane of these autophagosomes with lysosome (vacuoles in yeast and plants) exposes the inner membrane and engulfed material to acidic hydrolases that degrade the content. Autophagy can be divided into selective and non-selective processes. These are differentially regulated depending on the substrate and cellular conditions. While there is overlap amongst proteasome and autophagy substrates, autophagy is uniquely capable of clearing protein aggregates, damaged organelles, viruses, and large multi-subunit complexes (Ogata *et al*., 2006; Kraft, Reggiori and Peter, 2009; Kamada *et al*., 2010; Deffieu *et al*., 2013; Marshall *et al*., 2015; Mochida *et al*., 2015; Waite *et al*., 2015; BAO *et al*., 2016).

Proteasomes are one of the large multi-subunit complexes that are degraded via autophagy, a process referred to as proteaphagy. Proteaphagy appears to be a selective process as it depends on factors not involved in general autophagy (Waite *et al*., 2015; Cohen-Kaplan *et al*., 2016; Marshall, McLoughlin and Vierstra, 2016; Nemec *et al*., 2017; Li *et al*., 2019) and does not occur under several conditions that induce general autophagy (Waite *et al*., 2021). A striking example of this is carbon starvation, which induces general autophagy (Adachi, Koizumi and Ohsumi, 2017). Proteasomes, however, localize to cytosolic punctate structures termed proteasome storage granules (PSGs) upon carbon starvation (Laporte *et al*., 2008). While several factors that regulate PSG formation or their subsequent dissolution have been identified (Peters Lee Zeev *et al*., 2013; Weberruss *et al*., 2013; van Deventer *et al*., 2014; Marshall and Vierstra, 2018; Li *et al*., 2019), the signals that trigger proteasome relocalization and the mechanisms that regulate it have not been resolved.

Grown in the presence of glucose, yeast mainly utilize glycolysis for energy (ATP) and biomass production, while suppressing mitochondrial respiration (Kayikci and Nielsen, 2015). However, upon depletion of glucose, cells adapt and the metabolism switches from mainly glycolysis to mitochondrial respiration. This process is facilitated by autophagy, which provides a source of serine required for mitochondrial one-carbon metabolism (May, Prescott and Ohsumi, 2020). While uniquely adapted to utilize glucose as a carbon source, yeast grown in the presence of other carbon sources, like raffinose or glycerol, primarily utilize mitochondrial respiration for energy production (Kayikci and Nielsen, 2015; Adachi, Koizumi and Ohsumi, 2017). Interestingly, yeast do not induce general autophagy when switched from glucose containing media to carbon starvation media, presumably due to the lack of ATP. However, yeast grown on non-glucose carbon sources prior to starvation do induce general autophagy upon removal of the carbon source (Adachi, Koizumi and Ohsumi, 2017). This suggests the cellular respiratory state and ATP levels are determinants in signaling autophagy. However, it should be noted that the difference between glycolytic growth with repressed respiration and growth with active mitochondrial respiration is also accompanied by a differences in the expression of numerous genes (Galdieri *et al*., 2010).

As proteasomes and autophagy both contribute to the replenishment of cellular metabolites, we hypothesized that the metabolic state of cells, like with autophagy, impacts proteasomes as well. Here, we report the relocalization of proteasomes to PSGs is restricted when yeast are starved following respiratory growth, but not glycolytic growth (where respiration is suppressed). Consistent with this, conditions that interfered with mitochondrial function, such as inhibition of different electron transport chain complexes, resulted in the formation of proteasome granules in yeast. This differential regulation based on the initial carbon source involves specific signaling pathways: the AMP kinase Snf1 and a MAP kinase signaling cascade, both regulate proteasome granule formation. We propose a model where proteasome localization is regulated through mitochondrial respiration.

## Results

### Proteasome Granule Formation is Restricted During Respiratory Growth

The observation that the growth media prior to carbon starvation influenced yeast’s autophagic response resolved a controversy regarding general autophagy induction following carbon starvation (Takeshige *et al*., 1992; Lang *et al*., 2014). Starvation following respiratory growth (like in media containing raffinose or glycerol as the carbon source), promoted autophagy induction while growth in glycolytic/fermenting media, like dextrose (D-glucose), did not (Adachi, Koizumi and Ohsumi, 2017). Consistent with this, a link between respiration and autophagy induction during energy deprivation has been established (Yi *et al*., 2017).Thus, the metabolic state of the cell is an important variable when evaluating carbon starvation responses. We previously showed that carbon starvation does not induce proteasome autophagy in yeast (Waite *et al*., 2016), however we did not specifically control for the pre-starvation condition of the cells. Therefore, we used media containing dextrose (YPD, glycolysis), raffinose (YPR, glycolysis and respiration), or glycerol (YPG, respiration exclusively) as sources of carbon to examine proteaphagy following carbon starvation. We monitored proteasome autophagy with a GFP-tag fused to the regulatory particle subunit Rpn1. Here, the observation of vacuolar fluorescence in cells or the appearance of a faster migrating “free” GFP species on immunoblots are indicative of proteaphagy. No robust proteaphagy was observed upon switching to carbon starvation, independent of the growth medium prior to starvation. The amount of “free” GFP detected at 1 and 2 days of carbon starvation was less than “free” GFP detected following six hours of nitrogen starvation (Waite *et al*., 2015) (Fig. 1A). Instead, proteasome granules were induced, as previously reported (Waite *et al*., 2015; Marshall and Vierstra, 2018). Interestingly, pre-growth in dextrose resulted in approximately 58% of cells showing granules with 27% of these showing multiple granules per cell, whereas pre-growth in raffinose resulted in less than 32% of cells showing granules and 5% of these showing multiple granules per cell (Fig. 1B). A similar trend was observed when we monitored the core particle using a1-GFP as a reporter (S1). This suggest that proteasome localization is affected by the carbon source cells were grown in prior to starvation.

**Figure 1.**
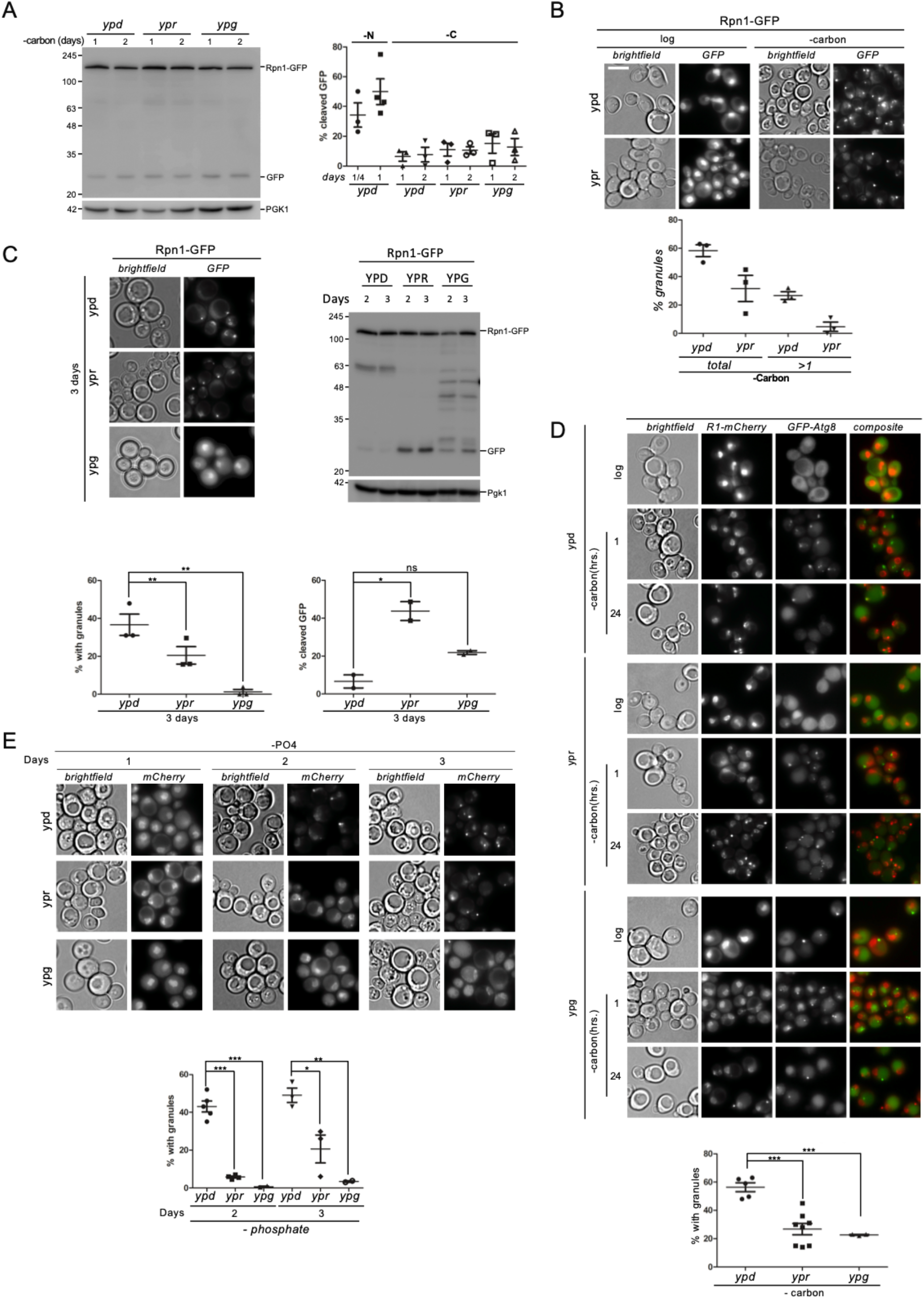
Proteasome granule formation is restricted during respiratory growth. **(A)** Rpn1-GFP expressing cells were grown for four hours in rich media containing dextrose (ypd), raffinose (ypr) or glycerol (ypg), followed by growth in SD media lacking carbon. 2 ODs of cells (i.e., the cells equivalent to two ml of culture OD_600_ 1) were harvested at indicated times and lysed as described in materials and methods. Lysates were separated by SDS-PAGE and immunoblotted for GFP and Pgk1. Data presented are representative of three independent experiments. Quantifications show the percentage of free GFP relative to the total amount of GFP (i.e., unprocessed + free GFP) observed following nitrogen or carbon starvation. **(B)** Cells grown as in A were imaged at log phase and following 24 hours carbon starvation. Quantification shows the percentage of cells that induced proteasome granules and the percent of those cells that formed two or more granules (Data presented are representative of three independent experiments with n>100 for each). Scale bar represents 5 μm. **(C)** (left) Rpn1-GFP expressing cells were grown for 3 days in the indicated media then imaged microscopically. Quantification shows the percent of cells with granules from three independent experiments. Statistical significance was determined with paired t-test (left) and n>100 for each datapoint. (right) 2 ODs of cells were collected from cultures as in the left panel and lysed for immune blotting against GFP or Pgk1. Data presented are representative of two independent experiments. Quantification shows the percentage of free GFP, which indicating proteasome autophagy, relative to total GFP. Significance was determined using an unpaired t-test. **(D)** Rpn1-mCherry (R1-mCherry); GFP-Atg8 expressing cells were grown in rich dextrose (ypd), raffinose (ypr) or glycerol (ypg) media followed by a change to carbon starvation media. Microscopy was performed at indicated times and quantifications show the percentage of cells with granules after 24 hours. Unpaired t-test was used to determine significance and n>100 for each data point. **(E)** Rpn1-mCherry expressing yeast were grown in indicated rich media for 4 hours, washed and switched to phosphate starvation media. Microscopy was performed at the indicated times. Quantification shows the percent of cells that formed granules at 2 and 3 days of phosphate starvation. Unpaired t-test was used to determine significance with n>100 for each datapoint.

To explore the role of nutrients in proteasome localization further, we analyzed the induction of PSGs following prolonged growth in rich media, another condition that results in PSG formation (Laporte *et al*., 2008; Peters *et al*., 2016). When granules formed after growth for an extended period, a single dominant granule was observed in cells regardless of the initial carbon source. However, we observed that cells grown in rich media containing dextrose for 3 days had more cells forming granules in general than cells grown in media containing raffinose. Growth in glycerol media resulted in very few cells forming granules after 3 days (Fig. 1C). This was also observed with α1-GFP as a reporter (S2). Consistent with a model where PSGs protect proteasomes from autophagy (Marshall and Vierstra, 2018), the conditions with less granules (i.e. raffinose and glycerol) showed more “free” GFP on immunoblots at both 2 and 3 days of growth in carbon containing media, (Fig. 1C, S3) suggesting an increase in proteaphagy. However, we would expect more proteaphagy following growth in glycerol considering this condition formed markedly fewer PSGs (Fig. 1C). Interestingly, we observed GFP-positive bands on immunoblots with molecular weights between “free GFP” and Rpn1-GFP following prolonged growth in glycerol. Such fragmentation patterns were recently observed specifically for GFP-tagged proteasomes degraded through ESCRT-mediated micro-autophagy (Li *et al*., 2019). In all, our data suggest that not only PSGs, but also the type and magnitude of proteasome autophagy, is dependent on the initial carbon source and the cell’s metabolic state prior to starvation.

Switching from dextrose media to carbon starvation results in the formation of multiple phagophore assembly sites (PAS), which does not occur when carbon starvation is initiated following prior growth in other carbon sources or upon nitrogen starvation. This likely affects autophagy induction and might contribute to the lack of autophagy in this condition (Adachi, Koizumi and Ohsumi, 2017). The number of proteasome granules observed per cell correlates with increased PAS sites when comparing switching from dextrose or raffinose to carbon starvation (Fig. 1B, S1). To determine if proteasomes co-localized with PAS during carbon starvation, we used cells expressing GFP-Atg8 (PAS/ autophagy marker) and Rpn1-mCherry. We monitored GFP and mCherry localization upon carbon starvation after pre-growth in dextrose, raffinose, or glycerol containing media (Fig. 1D). PAS structures were observed within 1 hour of starvation, when no PSGs are yet detectable. At 24 hours when both PSG and PAS can be observed, we did not observe any striking colocalization (Fig. 1D), indicating these are distinct structures. To test if PSG formation depended on PAS formation or the signaling pathway that induces an autophagic response upon carbon starvation, we analyzed strains deleted for different genes required for autophagy induction: ATG1, SNF1 and GGC1. These gene products activate Mec1 in a process that tethers Mec1 to the mitochondrial outer membrane upon carbon starvation. This mitochondrial localization together with active mitochondrial respiration are prerequisites for autophagy induction (Yi *et al*., 2017). ATG1, SNF1, and GGC1 were not required for proteasome granule formation upon carbon starvation (S4) indicating multiple distinct signals couple the cells metabolic state to these two degradative machineries. The majority of cells formed PSGs at 24 hours when pre-grown in dextrose (~56%), which was significantly higher compared to cells pre-grown in raffinose or glycerol (26% and 22% respectively) (Fig. 1D). In sum, conditions where cells initially had active respiration show less PSGs.

To further probe the role of mitochondrial respiration in regulating proteasome localization, we tested other conditions that induce respiration vs glycolysis. First, we tested the carbon starvation response after growing cells for 4 hours (more glycolysis and fermentation) or 24 hours (more respiration) in YPD medium. After 24 hours of dextrose depletion, yeast have switched to respiration for energy production (Galdieri *et al*., 2010). Indeed, starving after 24 hours of growth in dextrose resulted in a reduction in granule formation compared to 4 hours (S5). Next, we starved cells for phosphate following growth in different carbon sources. This was based on the rationale that cells lacking phosphate sources would be compromised in maintaining energy production as phosphate is required for ATP production (Ko, Hong and Pedersen, 1999). Phosphate starvation induces cell cycle arrest and a stress response similar to carbon starvation (Petti *et al*., 2011; Secco *et al*., 2012). Phosphate starvation also induced proteasome autophagy but to a lesser extent than nitrogen starvation at 24 hours (Waite *et al*., 2021). Here, we show that prolonged phosphate starvation induced proteasome granules (Fig.1E). Like PSGs induced by carbon starvation, these granules had different properties depending on the initial carbon source. Cells grown in raffinose that were starved for phosphate showed less PSGs compared to cells grown in dextrose. Even more striking was the almost complete absence of granules when cells were grown in glycerol (Fig. 1E), even though our phosphate starvation media always contained dextrose as a carbon source. Catabolite repression does not appear to play a role here as raffinose and glycerol pre-growth resulted in distinct phenotypes. Because granules induced by phosphate starvation are evident only at later timepoints (2 days), they could result from prolonged growth similar to granules formed during stationary phase. However, we found granules were induced to a much larger extent in SD media lacking phosphate compared to SD complete media, both of which contained dextrose as a carbon source (S6). In all, proteasome granule formation appears to be in part dependent on mitochondrial respiration. Under conditions of limited mitochondrial respiration, proteasomes appear to re-localize to PSGs more efficiently.

### Mitochondrial Inhibition Induces Proteasome Granule Formation

Cells grown in respiratory media formed fewer PSGs when starved for carbon, phosphate, or grown for three days. Thus, increased mitochondrial function appears to limit PSG formation. This led to the hypothesis that reducing mitochondrial function has opposing effect and causes proteasome granule formation. To test this, we inhibited mitochondrial function by targeting different components of the electron transport chain (ETC). Sodium azide, antimycin A, and oligomycin A were used to target complex IV, complex III, and ATP synthase respectively. We further utilized the uncoupler CCCP to disrupt ATP synthesis (Heytler and Prichard, 1962; Wikström and Berden, 1972; Gribble *et al*., 1997; Symersky *et al*., 2012). All four drugs induced proteasome granules in rich (dextrose) media (Fig. 2A), further corroborating the link between mitochondrial function and proteasome localization. To test if granules induced through mitochondrial inhibition behave like proteasome granules induced by carbon starvation, we tested the reversibility of these granules. Carbon starvation induced PSGs, unlike other types of proteasome containing granules, quickly disappear with proteasomes re-localizing to cell nuclei upon re-addition of a carbon source (Laporte *et al*., 2008; Peters Lee Zeev *et al*., 2013; Weberruss *et al*., 2013; Waite, Burris and Roelofs, 2020). Granules induced by mitochondria inhibitors quickly dissipated and GFP signal was predominantly nuclear after cells were washed and re-inoculated into drug free media (Fig. 2B, S7). This indicates that these granules are not associated with irreversible aggregates but are dynamic structures consistent with the proposed storage function of PSGs. Similar to ETC inhibitors, we predicted the absence of the final electron acceptor, oxygen, should also result in PSG formation by inhibiting oxidative phosphorylation. Indeed, upon growth in anoxic conditions, proteasome granules were induced (S8). Like with carbon starvation, the magnitude of mitochondria inhibition-induced granules was dependent on the carbon source during initial growth with the exception of Rpn1-GFP cells treated with antimycin A (Fig. 2C). These data further indicate a role for mitochondrial respiration in regulating proteasome granule formation.

**Figure 2.**
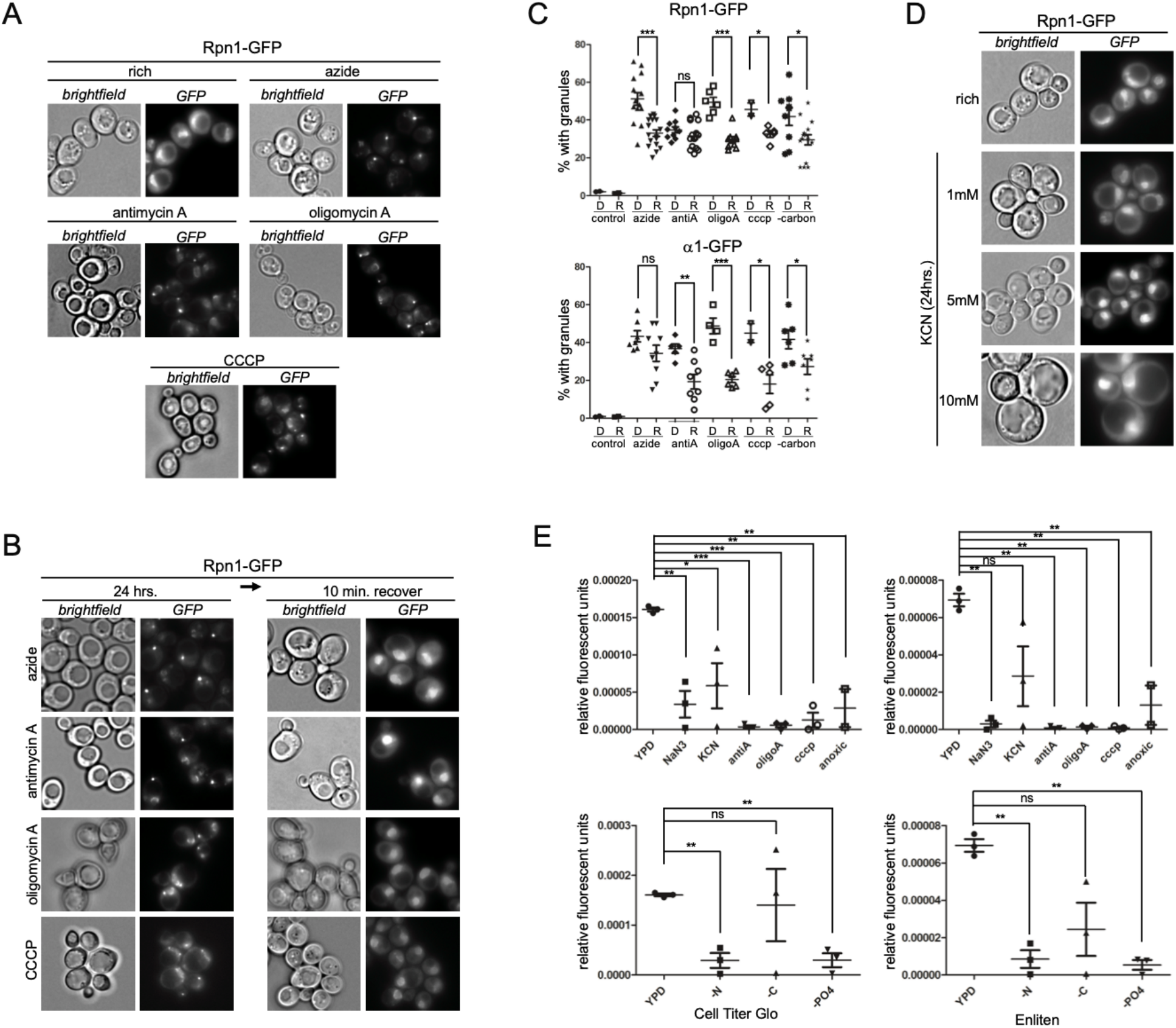
Mitochondrial inhibition induces proteasome granule formation. **(A)** Rpn1-GFP expressing cells were grown in rich media containing dextrose and treated with mitochondria inhibitors as described in materials and methods. Microscopy was performed 24 hours after inhibitor addition. Data are representative of three independent experiments. **(B)** Rpn1-GFP expressing cells treated for 24 hours with mitochondria inhibitors were washed and incubated in drug free media for 10 minutes. Data are representative of three independent experiments. **(C)** Rpn1-GFP or a1-GFP expressing cells were grown in media containing dextrose (D) or raffinose (R) for 4 hours, treated with mitochondria inhibitors, followed by and incubation for 24 hours. Microscopy was carried out and quantifications show the percentage of cells with granules. Unpaired t-tests were used to determine significance with n>100 for each datapoint. **(D)** Yeast expressing Rpn1-GFP were treated with increasing concentrations of potassium cyanide for 24 hours followed by microscopic analysis. Data are representative of three independent experiments. **(E)** ATP measurements were carried out following mitochondrial inhibition (upper panels) or nutrient starvation (lower panels) as described in materials and methods. Quantifications show relative ATP compared to untreated controls averaged from three independent experiments. Paired t-tests were used to evaluate significance.

Intriguingly, we did not detect induction of proteasome granules with all mitochondria inhibitors tested. Surprisingly, potassium cyanide (KCN) treatment, which like azide targets cytochrome c oxidase, did not result in any significant granule formation at various concentrations (Fig. 2D). The KCN treated cells grew slower than untreated cells (with a growth rate comparable to other mitochondria inhibitors) and we observed an increase in cell size at high concentrations (Fig. 2D), indicating KCN treatment was effective. Apparently mitochondrial inhibition or stress alone is not sufficient for PSG formation.

Mitochondria play a key role in efficient ATP production and ATP is essential for proteasome function (Finley, 2009; Schrader, Harstad and Matouschek, 2009). Furthermore, proteasome complexes are unstable *in vitro* in the absence of ATP (Kleijnen *et al*., 2007). Therefore we wondered if ATP maintenance and proteasome stability are key determinants in PSG formation; something that has been postulated before (Enenkel, 2018; Karmon and Ben Aroya, 2020). To test if PSG formation was linked to ATP production, we measured ATP under PSG inducing and non-inducing stimuli. To do this, we utilized the Cell Titer Glo as well as the Enliten ATP assay systems (Nicastro *et al*., 2015; Adachi, Koizumi and Ohsumi, 2017). Inhibiting mitochondria with sodium azide, antimycin A, oligomycin A, CCCP and growing yeast anoxically, led to a large reduction in ATP compared to growth in rich media (Fig. 2E). This reduction in ATP levels correlated with PSG formation (Fig. 2E). Interestingly, KCN treatment caused a smaller reduction in ATP levels compared to the untreated control, and this inhibitor did not induce PSGs.

We next measured ATP under starvation conditions. We observed a larger reduction in ATP following nitrogen starvation than carbon starvation, while only carbon starvation induced granule formation. Similarly, we observed a strong reduction in ATP detected following 24 hours of phosphate starvation, a condition that induces proteasome autophagy with PSGs observed 48 hours post starvation (Fig. 1F) (Waite *et al*., 2021).Thus, with starvation, a reduction in ATP levels by itself does not trigger proteasomes re-localization to PSGs. One possible explanation is that in addition to a drop in ATP levels, other signals are generated during nitrogen and phosphate starvation that result in proteaphagy.

### MAP Kinase Signaling is Required for Proteasome Granule Formation

The differential regulation of proteasome granules observed following respiratory vs non-respiratory growth, as well as the observation that not all mitochondria inhibitors induce proteasome granules (Fig. 2D), suggests that there is some regulator of this process that is only active under specific conditions. The cell wall integrity MAP kinase cascade proteins Mpk1(Slt2), Mkk1, and Mkk2 are known to regulate proteasome abundance and proteasome autophagy (Rousseau and Bertolotti, 2016; Waite *et al*., 2021). This kinase cascade has further been shown to be required for induction of general autophagy when antimycin A or potassium cyanide were added to cells in a process that was dependent on Atg32 and Atg11 (Deffieu *et al*., 2013). When we tested if deletion of *ATG32* disrupted the formation of proteasome granules induced after prolonged yeast growth, azide or antimycin A treatment, we observed no reduction in PSG formation (S9). Similarly, *ATG11* was not required for granule formation when cells were treated with sodium azide (S10). This shows that the requirements for autophagy induction and PSG formation are not identical in these conditions. Next, we wanted to test the role of MAP kinase signaling itself. To test if the MAPK Mpk1 was important for PSG formation, we deleted the MPK1 gene. In these cells we observed a striking difference depending on the initial carbon source used. PSG formation was unaffected in a *mpk1*Δ strain when starved for carbon following initial growth in dextrose (Fig. 3A, E). However, when yeast were grown in raffinose prior to starvation, Mpk1 was critical to form granules efficiently for both the RP (Rpn1) and CP (α1) reporters (Fig. 3A, E). This suggest that Mpk1 was required for proteasome granule formation in cells that had active mitochondrial respiration. When we monitored granule formation following oligomycin A, antimycin A and CCCP treatment, we found a similar requirement for Mpk1 when cells were grown in raffinose (Fig. 3B-D, F). With respect to sodium azide, granules monitored using a GFP-tag on the CP subunit α1 behaved as above with a reduction in the MPK1 mutant following growth in raffinose (Fig. 3E, F). Granule formation monitored using GFP-tag on the RP subunit Rpn1, however, was restricted when *mpk1Δ* cells were grown in dextrose but not raffinose (Fig. 3E, F). This observation further supports the idea of differential regulation of CP and RP in granule formation, as has been observed before (Weberruss *et al*., 2013; Marshall and Vierstra, 2018; Karmon and Ben Aroya, 2020). Next, we tested for a role of the proteasome chaperone Adc17. Adc17 is activated under several stress conditions (Hanssum *et al*., 2014) by Mpk1 (Rousseau and Bertolotti, 2016). However, Adc17 was dispensable for PSG formation, suggesting an alternate route of proteasome regulation by Mpk1(S11).

**Figure 3.**
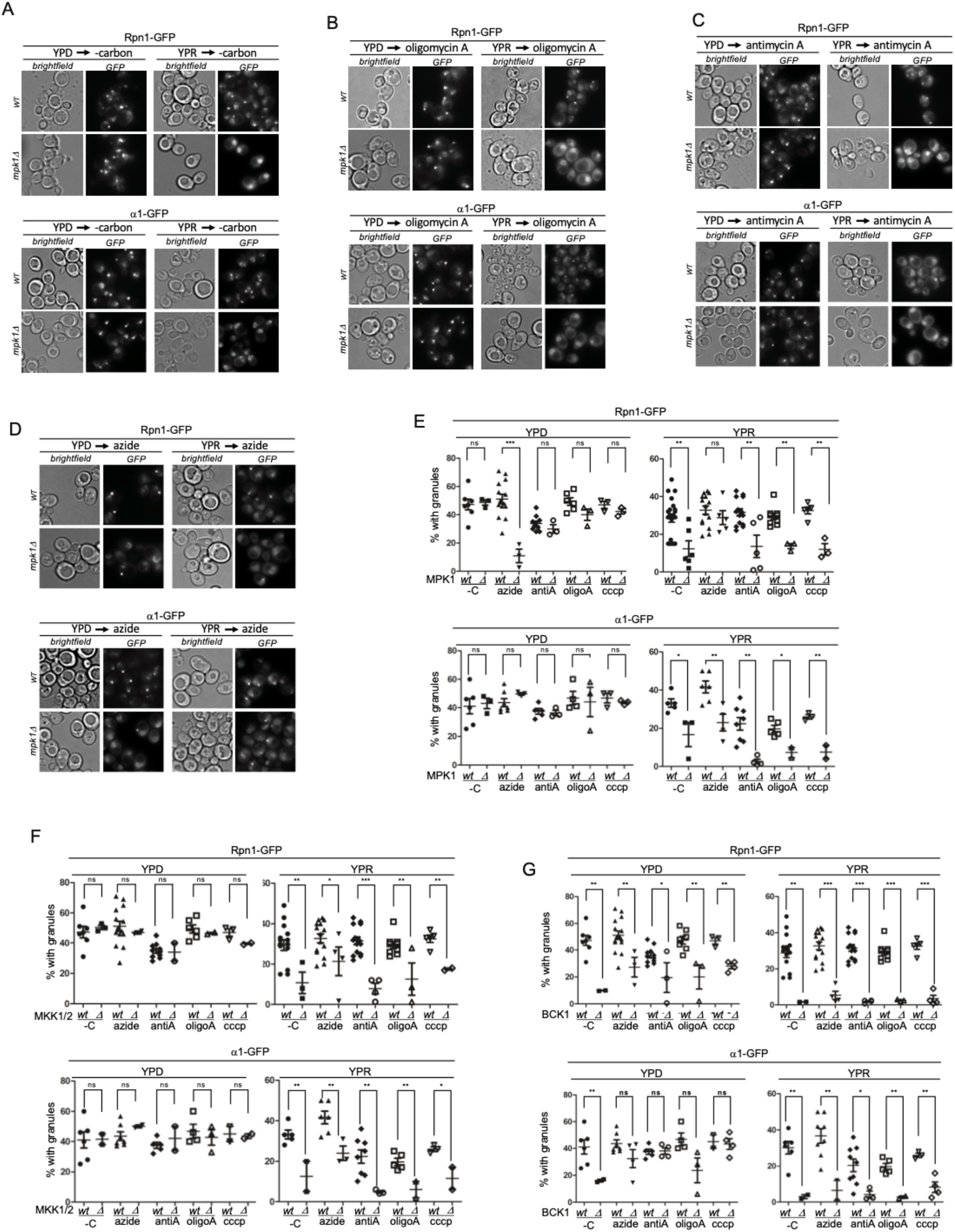
MAP kinase signaling is required for proteasome granule formation. **(A-D)** WT and *mpk1Δ* yeast expressing Rpn1-GFP or α1-GFP were grown to log phase in rich media containing dextrose or raffinose. Next, cells were starved for carbon, or treated with oligomycin A, antimycin A, or sodium azide for 24 hours and imaged. **(E)** Quantification of the percent of granule formation from yeast cultured and treated as in A-D, or CCCP. Statistical significance was determined using unpaired t-tests. Three or more independent experiments were quantified with n>100 for each datapoint. **(F)** wild type (wt) and MKK1 MKK2 double deletion yeast (Δ) expressing Rpn1-GFP or a1-GFP were grown in dextrose or raffinose for four hours, starved for carbon or treated with mitochondria inhibitors as above. Graphs show the percent of cells with granules after 24 hours with significance determined by unpaired t-tests. At least three independent experiments were quantified with n>100 for each datapoint. **(G)** wild type and *bck1Δ* yeast expressing Rpn1-GFP or α1-GFP were grown to log phase in rich media containing dextrose or raffinose then starved for carbon or treated with mitochondria inhibitors as above. Quantifications show the percent of cells that formed granules after 24 hours. Unpaired t-tests were used to determine significance. At least three independent experiments were quantified with n>100 for each datapoint.

To determine the involvement of other kinases, we analyzed a number of proteins within the cell integrity pathway, as well as other major yeast MAPKs from independent pathways (Levin, 2005). The deletion of *FUS3* or *HOG1* (MAPKs involved in pheromone response and osmoregulation respectively) did not prevent or substantially reduce proteasome granule formation (S11). This indicates a specific role for the cell integrity kinase pathway. Looking at MAP kinase kinases upstream of Mpk1, we evaluated Mkk1 and Mkk2. While individual deletion of these paralogs had little impact on granule formation, a double deletion of *MKK1* and *MKK2* showed only few proteasome granules under carbon starvation, azide treatment, antimycin A, oligomycin A and CCCP treatment (Fig. 3F).This reduction was, like with Mpk1, observed when cells were grown in raffinose but not glucose prior to treatment. Tracing this MAP kinase cascade further, we found that the MAPKKK Bck1 was similarly required, however these cells formed more granules after switching from glucose media than MPK1 or MKK1/MKK2 mutants. Though we did observe a more significant reduction compared to wildtype when cells were cultured in raffinose (Fig. 3G). Upstream of Bck1, the transmembrane activator of the cascade, Wsc1, was also required for granule formation (S12). Other upstream activators of the cell integrity pathway, namely Wsc2, Wsc3 and Ack1 (Verna *et al*., 1997; Kuranda *et al*., 2006; Krause, Xu and Gray, 2008), were not required for PSG formation under the same conditions (S12). Our data show that an intact map kinase pathway, starting at Wsc1 and moving downstream to Mpk1, is required for proteasome granule formation when cells are grown in respiration inducing conditions. We have previously reported that Mpk1, Mkk1 and Mkk2 are required for efficient proteaphagy (Waite et al, 2021). Both conditions require nuclear export of proteasomes, potentially indicating this kinase cascade regulates nuclear export of proteasomes, for example by Mpk1 directly phosphorylating proteasomes. However, Mpk1 is involved in many cellular processes and this kinase pathway might be indirectly involved in signaling proteasome re-localization. These possibilities are currently being tested.

### Snf1 is required for proteasome granule formation upon mitochondrial inhibition

As mentioned above, Snf1 is required for the induction of autophagy upon glucose starvation. This kinase is recruited to mitochondria shortly after the onset of glucose starvation where it phosphorylates Mec1. Phosphorylated Mec1 recruits Atg1 to mitochondria and both factors are required for maintaining mitochondrial respiration during glucose starvation (Yi *et al*., 2017). This respiration in turn is also required for autophagy induction (Adachi, Koizumi and Ohsumi, 2017; Yi *et al*., 2017). We did not observe a reduction in PSGs upon carbon starvation when SNF1 or other factors involved in this pathway were deleted (S4), a finding consistent with published data (Li *et al*., 2019). Snf1 was also not required for proteasome autophagy induced by nitrogen starvation, but was required for proteasome autophagy observed following 4 days of growth in SC media with 0.025% (low) glucose (Li *et al*., 2019). Intriguingly, we noticed a different role for Snf1 in proteasome autophagy under growth with different carbon sources. Deletion of SNF1 promoted autophagy of proteasomes when cells were grown for 3 days in rich media containing dextrose or glycerol but not raffinose (Fig. 4A). Further, we observed that SNF1 mutants failed to form proteasome granules efficiently under these conditions (Fig. 4B). This is consistent with a model whereby PSGs protect proteasome from autophagic degradation (Marshall and Vierstra, 2018), however, we did not observe increased autophagy when cells were grown in raffinose, despite the failure in granule formation in this condition. Here, GFP signal appeared more nuclear and diffuse in the cytoplasm for the Snf1 mutant than the wildtype which formed PSGs (Fig. 4B). The disparate phenotypes we observed likely reflect the complexity of combined inputs to cells metabolic state and mitochondrial respiration activity in regulating proteasome localization.

**Figure 4.**
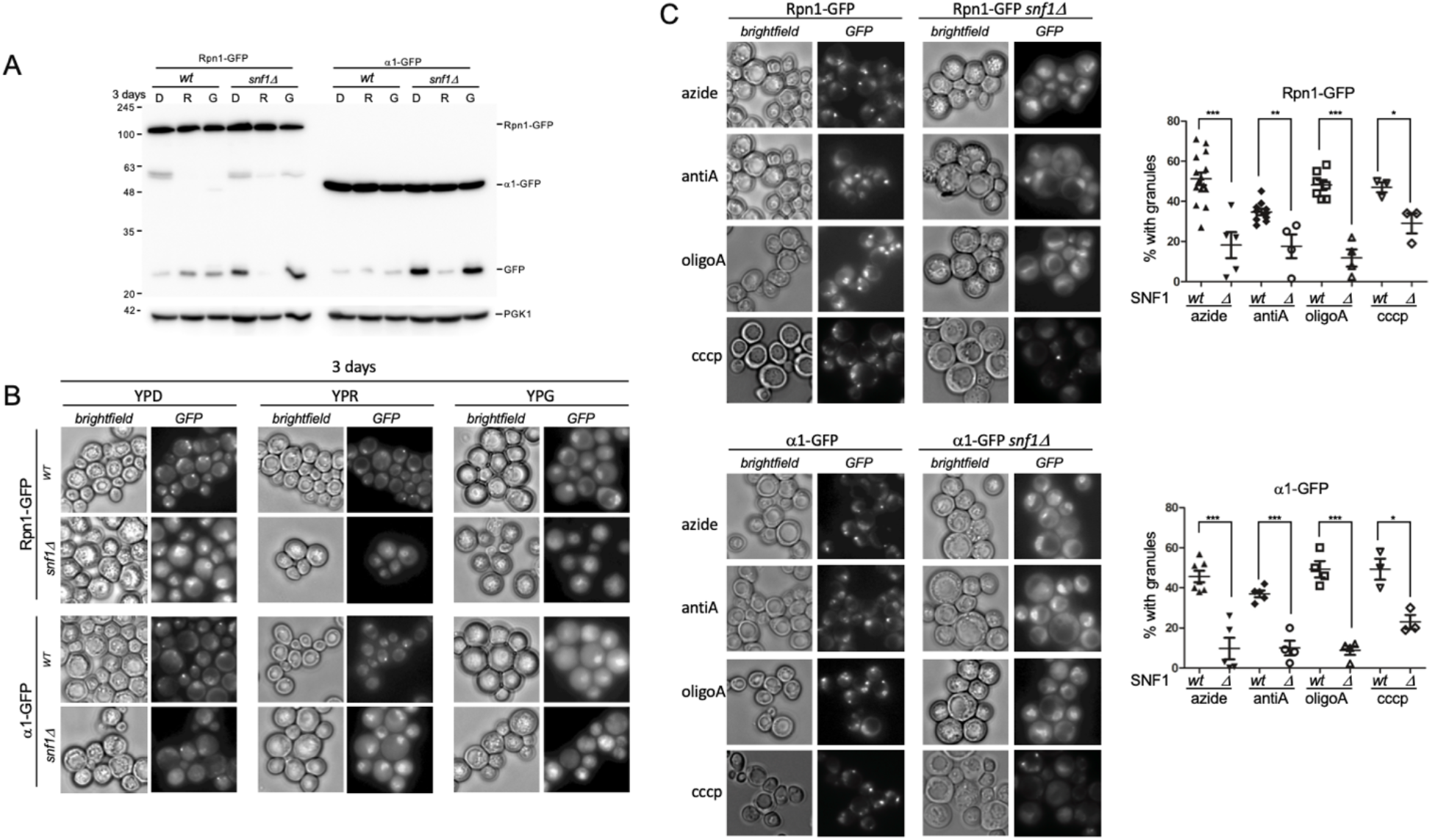
Snf1 is required for proteasome granule formation upon mitochondrial inhibition. **(A)** wild type (wt) and SNF1 deleted yeast expressing Rpn1-GFP or α1-GFP were grown for three days in rich media containing dextrose, raffinose or glycerol (D, R and G respectively). Cells were harvested and lysed as described in materials and methods. Immune blotting for GFP and Pgk1 was performed. Data are representative of two independent experiments. **(B)** Cells from A were imaged microscopically to observe proteasome localization. Data are representative of two independent experiments. **(C)** Yeast as in A and B were grown in YPD medium to log phase and starved for carbon or treated with mitochondria inhibitors. Microscopy was performed and quantifications show the percent of cells that form granules following mitochondria inhibition in the SNF1 deleted yeast compared to wild type. Significance was determined using unpaired t-tests. At least three independent experiments with n>100 for each datapoint, were used for quantification.

To gain more insight into the role of Snf1 in regulating proteasomes localization, autophagy, and granule formation, we tested the effect of mitochondria inhibitors in these cells. Unlike granules induced by carbon starvation, granules induced with mitochondria inhibitors were highly dependent of Snf1 (Fig. 4C) for both the RP and CP tag reporters (Li *et al*., 2019). Upon mitochondrial inhibition, compared to wild type, Snf1 mutants have fewer proteasome granules, more nuclear GFP signal and little to no vacuolar GFP signal. Of note, these cells were grown in glucose where Snf1 is inactive (Schüller, 2003; Kayikci and Nielsen, 2015). This may indicate an alternative role for Snf1 in regulating proteasome localization outside of its canonical role of non-fermentative gene repression. In all, these data suggest that different pathways are involved in the formation of proteasome storage granules, for example during carbon starvation versus mitochondria inhibition. This raises the question to what extent these granules are similar or qualitatively different in function or regulation.

## Discussion

The transcriptional upregulation of proteasomes (by Rpn4 in yeast and Nrf1 in mammals) is critical for cells to respond to various stress conditions. However, not all stresses require a (continuous) upregulation of proteasomes and instead trigger either the degradation of proteasomes or their relocalization into cytosolic granules. In yeast, proteasome complexes are enriched in the nucleus under optimal conditions such as during logarithmic growth. However, proteasomes re-localize to cytoplasmic granules upon carbon limitation (Enenkel, 2012; Peters Lee Zeev *et al*., 2013; Waite *et al*., 2015; Marshall and Vierstra, 2018). When stressed for nitrogen, on the other hand, proteaphagy is induced (Marshall *et al*., 2015; Waite *et al*., 2015; Nemec *et al*., 2017). Similarly, in human cells, proteasomes have been reported to undergo proteaphagy in response to amino acid starvation or upon proteasome inhibition (Cohen-Kaplan *et al*., 2016; Choi *et al*., 2020; Goebel *et al*., 2020). Interestingly, cytoplasmic proteasomes appear to be substrates for proteaphagy upon amino acid deprivation while nuclear proteasomes undergo liquid-liquid phase separation (LLPS) (Uriarte *et al*., 2021). Osmotic stress has also been shown to induce nuclear LLPS granules of proteasomes (Yasuda *et al*., 2020). Thus, both in yeast and humans depending on the condition, proteasomes show distinct fates and localization. To understand how these specific fates of proteasomes are regulated, we need to not only know the factors involved, but also understand the triggers responsible. This can be rather subtle, as it was recently shown that clearly defined fates for proteasomes were observed when comparing cells starved with low levels of glucose or no glucose at all (Li *et al*., 2019). Here, only the cells starved in low glucose induced Snf1-dependent micro-autophagy of proteasomes. In the current study, we sought to determine in detail what triggers proteasome granular localization and further identified some of the key signaling kinases involved.

## Mitochondrial Respiration as key determinant

Our data show that mitochondrial respiration, at least in part, regulates proteasome localization (Figs. 2, 5). First, we show that several inhibitors of mitochondrial respiration robustly induce PSG formation. Second, removing a carbon source from cells that grew in media that suppresses respiration (i.e. with glucose (Galdieri *et al*., 2010)) resulted in the formation of multiple granules per cell, whereas one dominant granule was present when cells were switched from media that required respiration (glycerol or raffinose). This observation shows interesting parallels with the regulation of general autophagy upon carbon starvation that was recently shown to be dependent on respiration (Adachi, Koizumi and Ohsumi, 2017). Another study found a complex of Snf1-Mec1-Atg1 is recruited to the mitochondrial membrane by Ggc1 following the onset of carbon starvation. The formation of this complex is required to maintain active mitochondrial respiration which in turn is required to initiate carbon starvation induced autophagy (Yi *et al*., 2017). Surprisingly, the deletion of GGC1 did not impact PSG formation, suggesting additional signaling pathways link respiration to the regulation of proteasome localization.

**Figure 5.**
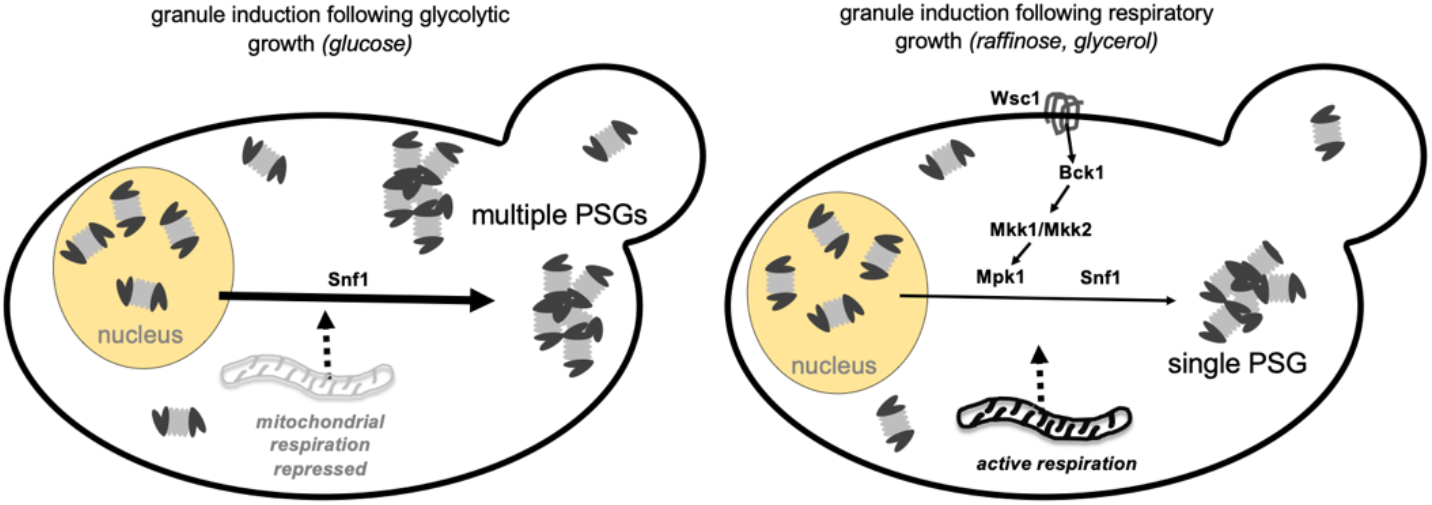
Model for PSG formation following glycolytic or respiratory growth. Switching from glycolytic growth media (i.e., with repressed mitochondrial respiration) to carbon starvation results in the formation of multiple proteasome granules per cell and more proteasome granules as compared to cells starved after growth in respiratory media. The kinase Snf1 is required for proteasome granule formation upon mitochondrial inhibition but not carbon starvation. The cell integrity MAP kinase cascade (from Wsc1 to Mpk1) is required from proteasome granule formation following respiratory but not glycolytic growth.

The extent of granule formation when cells were grown in different carbon sources negatively correlated with the amount of general autophagy induced, as we detected a reduction in the number of PSGs under conditions that led to increased general autophagy upon carbon starvation (Adachi, Koizumi and Ohsumi, 2017). Here, we observed only a small increase in proteaphagy (Fig. 1A) when cells were grown in respiratory media, suggesting the reduced levels of PSGs cannot be explained by increased autophagy of proteasomes. This is in contrast to the reported role of PSGs in protecting proteasomes from autophagy as we would expect more proteasome autophagy when there is a reduction in PSGs. Our data suggest that proteasomes are excluded from autophagic degradation without being sequestered into PSGs. This may indicate a role for proteasome activity in regulating the autophagic response of respiring cells. Further supporting our observations is the lack of an increase in proteasome autophagy under autophagic conditions in mutants that are defective in granule formation (Figs. 3 and 4). The nuclear enrichment of fluorescence we observed under these conditions indicates that the majority of proteasomes remained nuclear. This localization shields proteasomes from general autophagy (Waite *et al*., 2015; Nemec *et al*., 2017). Thus, proteasomes can be excluded from autophagic degradation either by PSG formation or by nuclear retention. In line with this, the deletion of autophagy related genes results in nuclear retention (Waite *et al*., 2015, 2021) This suggest proteasome relocalization is tightly regulated and below we discuss a kinase cascade we have identified that is involved in this process.

## Kinases that regulate proteasome localization

MAP kinases and Snf1 signaling pathways both regulate proteasome localization, however, each under distinct conditions. The Mpk1 kinase cascade was required following growth in media that induced respiration, while Snf1 was required upon mitochondrial inhibition (but not carbon starvation). These pathways are known to cooperate upon cell wall and oxidative stress (Backhaus *et al*., 2013; Willis *et al*., 2018), indicating they might work together to regulate proteasomes more generally. Indeed, both pathways are also involved in regulating proteasome abundance and mobilization upon different stressors. Mpk1 is required for proteasome upregulation upon ER stress (Rousseau and Bertolotti, 2016; Schmidt *et al*., 2019) as well as required for efficient proteaphagy (Waite *et al*., 2021), and Snf1 is required for micro-autophagy of proteasomes following growth in low glucose media (Li *et al*., 2019). The requirement for these pathways in both proteasome autophagy and proteasome granule formation could indicate they are necessary for efficient nuclear export of proteasomes, a pre-requisite for both autophagy and granule formation (Nemec *et al*., 2017). However, the possibility that they play a less direct role in regulating proteasome localization cannot be excluded. It has been shown that MPK1 is required for general autophagy induction when mitochondria are inhibited with the drug antimycin A (Deffieu *et al*., 2013). Similarly, it is required for mitophagy and pexophagy induced following nitrogen starvation, even though general autophagy does not depend on Mpk1 (Mao *et al*., 2011). Mitophagy here was shown to depend on the complete the cell integrity MAPK cascade (from the transmembrane receptor kinase Wsc1 to Mpk1). We demonstrate that this same cascade is required for proteasome granule formation. These data point to a more general signaling role for MPK1, particularly when mitochondria are affected. However, other factors that are required for mitophagy under this conditions, such as Atg32, and Atg11, had no effect on the formation of proteasome granules, suggesting that a general failure in mitophagy does not correlate with a failure in proteasome granule formation. Nevertheless, the MPK1 kinase cascade is clearly important in regulating proteasome localization, particularly following respiratory growth.

An intact MAP kinase cascade is not sufficient to promote proteasome granule formation. We report here that deletion of SNF1 restricted proteasome granule formation upon inhibition of mitochondria. It was previously shown that this kinase is not required for granule formation when cells were starved for glucose (Li *et al*., 2019). The requirement for this protein in these different context suggests that proteasome granules induced upon carbon starvation versus mitochondrial inhibition are distinct. It is intriguing that Snf1 was required for granule formation when cells were grown in glucose rich media, as this protein is known to be inactive in this condition (Schüller, 2003; Turcotte *et al*., 2010). This suggests an unidentified role for Snf1 in this context that regulates proteasome localization.

In all our data show that proteasomes are specifically regulated under different metabolic conditions. This may indicate that proteasome activity is similarly regulated based on cellular needs. Given that the majority of proteasomes are nuclear, proteasome capacity and activity must be altered when they re-localize. Indeed, the current literature suggests that proteasomes are inactive in PSGs (Gu *et al*., 2017; Enenkel, 2018). Although it should be noted that LLPS structures induced by osmotic stress in HCT116 colon cancer cell line appear to be actively degrading ribosomal proteins (Yasuda *et al*., 2020). Under conditions of mitochondrial respiration where proteasomes are more nuclear, proteasome activity may be necessary. This observed relationship between mitochondrial respiration and proteasome localization is intriguing, as other studies have demonstrated a functional link between mitochondria and proteasomes (Bragoszewski *et al*., 2013; Bragoszewski, Turek and Chacinska, 2017; Lavie *et al*., 2018).This is also evident in Parkinson’s disease where proteasomes play a crucial role in regulating mitochondrial dynamics (Junn *et al*., 2002; Ciechanover and Brundin, 2003; Webb *et al*., 2003; Um *et al*., 2010). Further, proteasomes are required to resolve mitochondrial stress induced upon transporter clogging or accumulation of misfolded mitochondrial proteins in the cytoplasm (Boos *et al*., 2019). Our data show that proteasomes are responsive to cellular metabolism and are regulated differently depending on the cell’s metabolic status. While ATP appears to not be the trigger for PSG formation, at least upon starvation, our data make it clear that mitochondrial function and signaling play an important role.

## Materials and Methods

### Yeast Strains

All strains used in this study are reported in table S1. Our background strains are the W303 derived SUB61 (Matα, *lys2-801 leu2-3, 2-112 ura3-52 his3-Δ200 trp1-1*) that arose from a dissection of DF5 (Finley, Özkaynak and Varshavsky, 1987). Standard PCR based procedures (primers and plasmids presented in table S2) were used to delete specific genes from the genome, or introduce sequences at the endogenous locus that resulted in the expression of C-terminal fusions of GFP or mCherry (Goldstein and McCusker, 1999; Hailey, Davis and Muller, 2002; Janke *et al*., 2004).

### Yeast Growth Conditions

Overnight cultures of yeast were inoculated at an OD_600_ of 0.5 and grown in yeast extract peptone (YEP) medium supplemented with 2% dextrose, raffinose or glycerol as a carbon source and grown to an OD_600_ 1.5 (approximately 4 hours). To induce starvation, cultures growing logarithmically were centrifuged, washed with respective starvation medium, re-inoculated at an OD_600_ of 1.5, and incubated at 30 °C with constant shaking. Yeast nitrogen base lacking carbon or phosphate sources was used to make the respective starvation media. For drug treatments, cultures were grown to an OD_600_ of 1.5 as above, then treated. Sodium azide (VWR), antimycin A (Sigma), oligomycin A (Cayman Chemical) and 2-[2-(3-chlorophenyl)hydrazinylidene]-propanedinitrile (CCCP) (Sigma), were used at final concentrations of 0.5 μM, 0.1 mM, 2.5 μM and 10 μM respectively. Potassium cyanide (Fisher) was used at concentrations shown in Fig. 2E. Anoxic growth was performed by transferring 1.8mL of logarithmically growing culture to a 2 mL culture tube that was sealed and incubated without shaking for 24 hours at 30 °C.

### Protein Lysates and Electrophoresis

For western blots, 2 ODs of cells were collected at indicated timepoints and treatments and stored at −80 °C. Lysis was completed using previously establish methods (Kushnirov, 2000). Following electrophoreses, samples were transferred to PVDF membranes and immuno-blotted for GFP then Pgk1 followed by the appropriate horseradish-peroxidase conjugated secondary antibodies. Antibodies used were anti-GFP (1:500; Roche, #11814460001), and anti-Pgk1 (1:10,000; Invitrogen, #459250). Horseradish-peroxidase activity was visualized using the Immobilon Forte Western HRP substrate (Millipore), and images were acquired using the G-box imaging system (Syngene) with GeneSnap software. Genetools was used to quantify the amount of free GFP normalized to the Pgk1 loading control.

### ATP Measurements

Enliten ATP assay: 1 OD of cells was collected following indicated treatments and frozen in liquid nitrogen. Pellets were resuspended in 50 μL of 2.5% trichloro-acetic-acid (TCA) and boiled for 3 mins. Sample was centrifuged (13,000 RPM) for 1 min and 2 μL supernatant was added to 98 μL of 25 mM Tris-HCL [pH 8.8] (1:50 dilution). 10 μL of 1:50 dilution was combined with 40 μL of 25 mM Tris-HCL [pH 8.8] in a white 96 well plate. 50 μL of rLuciferase/Luciferin reagent (Promega, ENLITEN^®^ ATP Assay System) was added to each well and luminescence was monitored using a plate reader.

Cell Titer Glo ATP assay: 0.5 OD of cells were centrifuged following indicated treatments and resuspended in 50 μL of sterile water. Samples were transferred to a 96 well black plate and 50 μL of Cell Titer Glo 2.0^®^ reagent was added. This plate was incubated in the dark under constant rotation on an orbital shaker for 4 minutes. Following rotation, the plate was further incubated in the dark for 10 minutes at room temperature. Luminescence was then measured using a plate reader.

### Fluorescence Microscopy

All microscopy was performed with live yeast where proteasome subunits Rpn1 or a1 was C-terminally tagged at their endogenous locus with expression driven by the endogenous promoter. GFP-Atg8 was produced as previously described (Li *et al*., 2015). After indicated treatments, approximately 2 ODs of cells were pelleted, washed with PBS, then resuspend in 30 μL of PBS. 3 μL of this sample was then mounted on 1% soft agar slides as described by E. Muller (https://www.youtube.com/watch?v=ZrZVbFg9NE8) (Sundin *et al*., 2004). All imaging by fluorescence microscopy was done within 10 mins following wash to avoid the effects of prolonged incubation on slides. Images were acquired at room temperature using a Nikon Eclipse TE2000-S microscope at 600X magnification with a Plan Apo 60x/1.40 objective equipped with a Retiga R3^tm^ camera (QImaging). Images were collected using Metamorph software (Molecular Devices) and analyzed using FIJI. All quantification was performed using FIJI.

## Acknowledgements

We thank Dr. Stella Lee for helpful discussions and feedback on the manuscript. We thank Mandeep Kaur for the generation of some of the yeast strains used in this study and feedback on the manuscript. We thank Dr. Alicia Burris for sharing her observation of proteasome granule formation under hypoxic conditions.

## Competing interests

The authors declare no competing or financial interests.

## Author contributions

Conceptualization: K.A.W., J.R.; Methodology: K.A.W., J.R.; Validation: K.A.W., J.R.; Formal analysis: K.A.W., J.R.; Investigation: K.A.W.; Writing - original draft: K.A.W.; Writing - review & editing: K.A.W., J.R.; Visualization: K.A.W., J.R.; Funding acquisition: J.R.

## Funding

This work was supported by grants from the National Institutes of Health and National Institute of General Medical Science (K-INBRE program P20GM103418 and R01GM118660 to J. R.). The content is solely the responsibility of the authors and does not necessarily represent the official views of the National Institutes of Health.

## Supplementary information

Supplementary information contains tables S1 and S2, supplementary figures S1-S12 and complete images of immunoblots used.

## Notes

### Competing Interest Statement

The authors have declared no competing interest.

