## supplement for "Proteasome granular localization is regulated through mitochondrial respiration and kinase signaling"

**Running title:** Mitochondrial function regulates proteasome localization

**Content:**

Supplementary figures S1-S12

Blot transparency: complete image of immunoblots.

Supplementary table 1. Yeast strains

Supplementary table 2. Primers

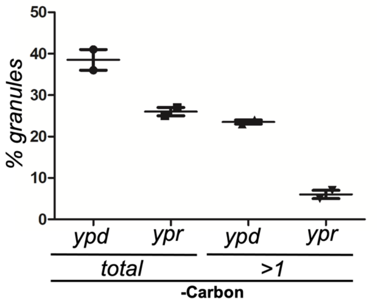


**S1.** α1-GFP expressing yeast were grown to log phase in rich media with dextrose or raffinose, switched to carbon starvation media and monitored for granule formation after 24 hours. Two independent experiments were quantified to determine the percent of cells that formed granules and the percent of granule forming cells that formed more than one granule. n>100 for each datapoint.

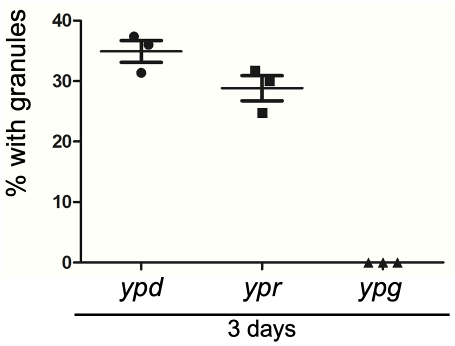


**S2.** α1-GFP expressing yeast were grown for three days in rich medium containing dextrose, raffinose or glycerol to compare magnitude of proteasome granule induction. Three independent experiments were quantified. n>100 for each datapoint.

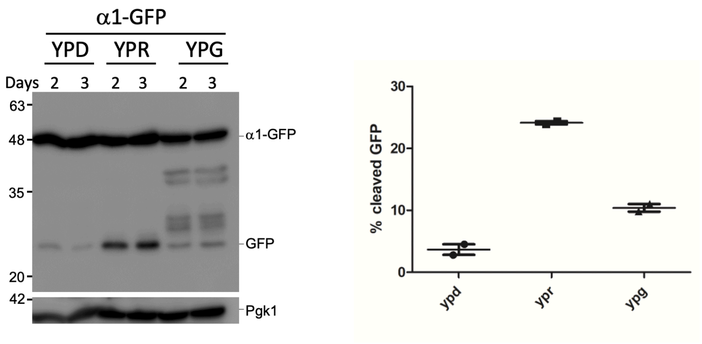


**S3.** α1-GFP expressing yeast were grown for two or three days in rich dextrose, raffinose or glycerol media. 2 ODs of cells were harvested, lysed, and analyzed by SDS-PAGE followed by immunoblotting against GFP or Pgk1. Data are representative of two independent experiments and quantification shows the percent cleaved GFP relative to total GFP.

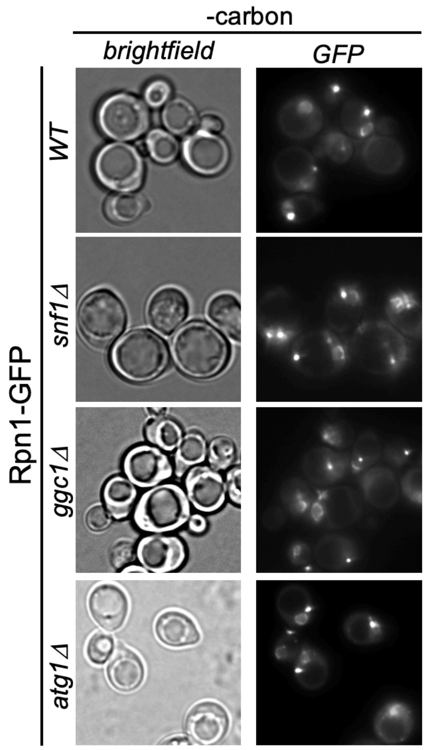


**S4.** Wild type (WT), *snf1Δ , ggc1Δ* and *atg1Δ* yeast expressing Rpn1-GFP were grown to log phase in rich media containing raffinose, followed by 24 hours incubation in carbon starvation media and microscopy analyses. Data presented are representative of three independent experiments.

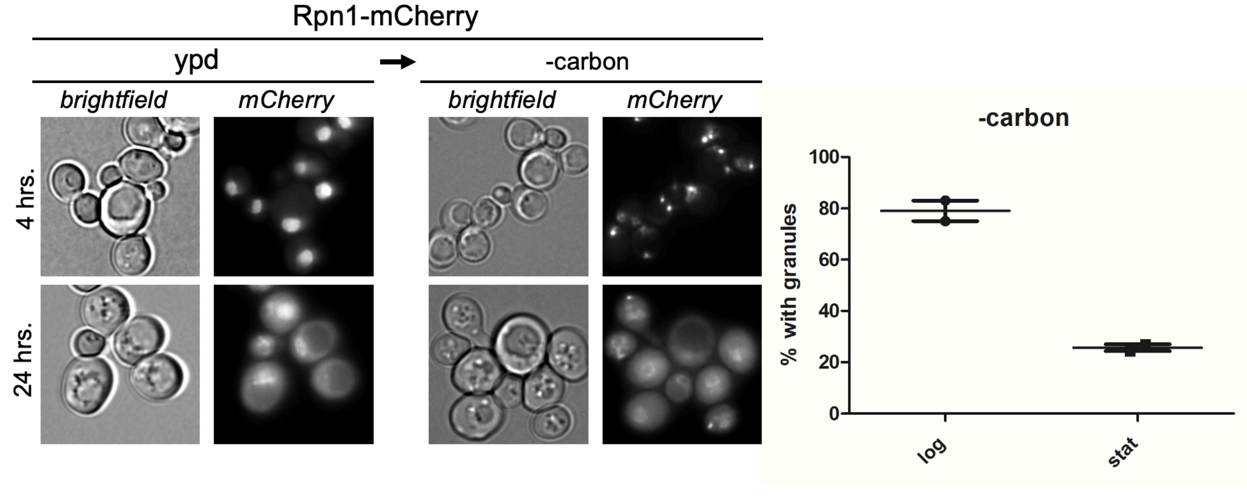


**S5.** Rpn1-mCherry cells were grown logarithmically (4 hours) or grown for 24 hours in rich dextrose media. Next, cells were switched to carbon starvation media and monitored after 24 hours. Quantifications show the percentage of cells with granules from two independent experiments. For each datapoint n>100.

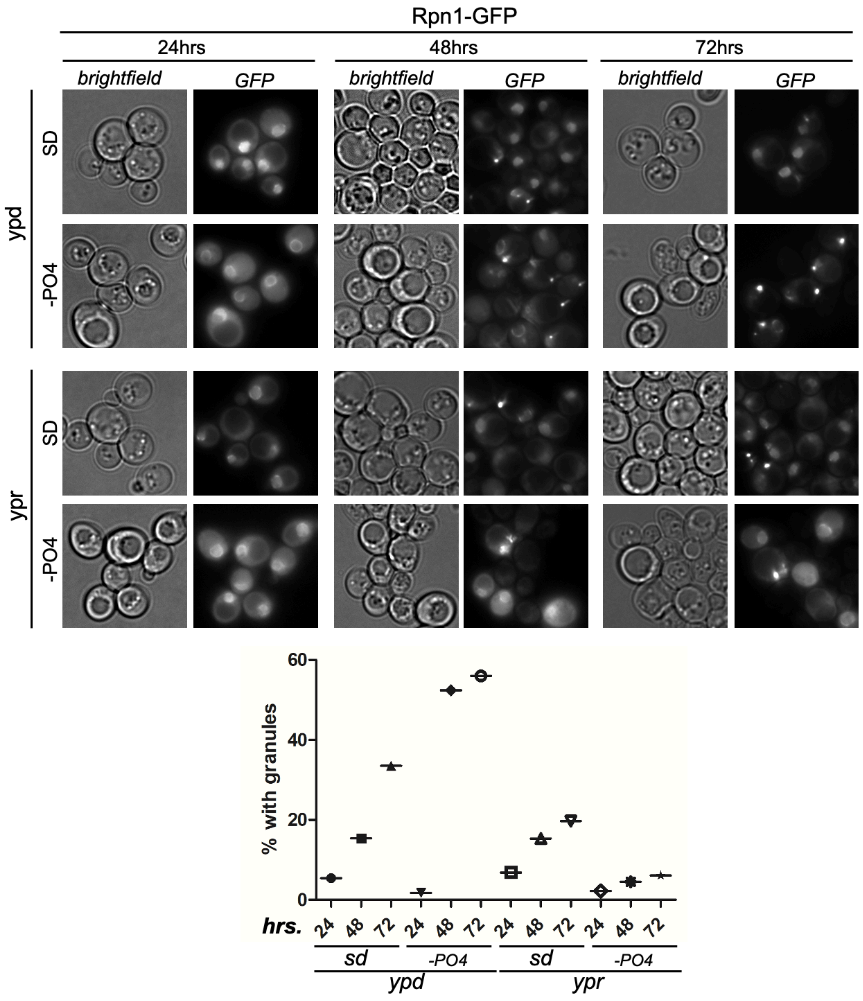


**S6**. Rpn1-GFP expressing cells were grown in rich dextrose or raffinose media for 4 hours and switched to SD complete media or SD media lacking phosphate. Microscopy was performed at indicated times and quantifications show the percentage of cells with granules from one experiment, which complements data presented in Fig. 1E. n>100 for each datapoint.

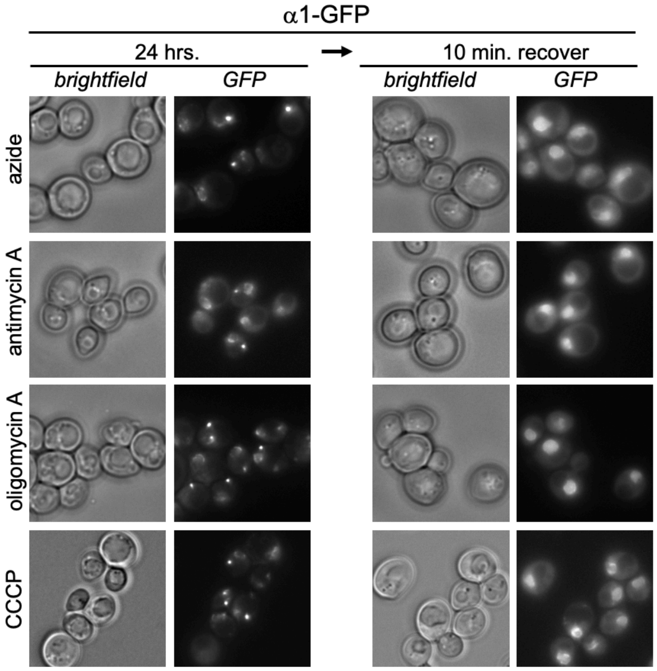


**S7.** α1-GFP expressing yeast were grown in rich dextrose media to log phase, treated with mitochondria inhibitors, and incubated for 24 hours followed by microscopy. At 24 hour incubation cells were also washed, incubated in drug free media for 10 minutes, and imaged. Data presented are representative of two independent experiments.

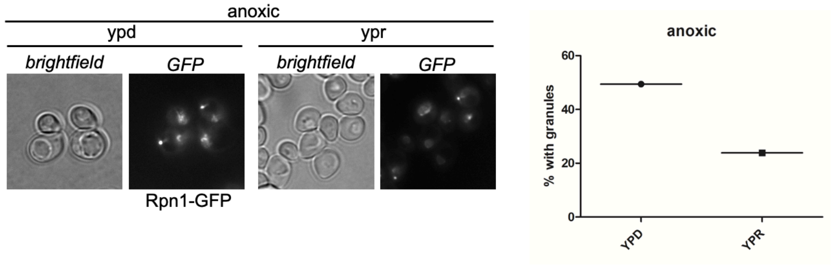


**S8**. Rpn1-GFP expressing cells were grown in media containing dextrose or raffinose for 4 hours, transferred to sealed culture tubes, incubated for 24 hours, and imaged. The induction of granules in hypoxic conditions has been observed in multiple independent experiments. Quantifications showing the percentage of cells with granules from one experiment and n>100 for each datapoint are shown.

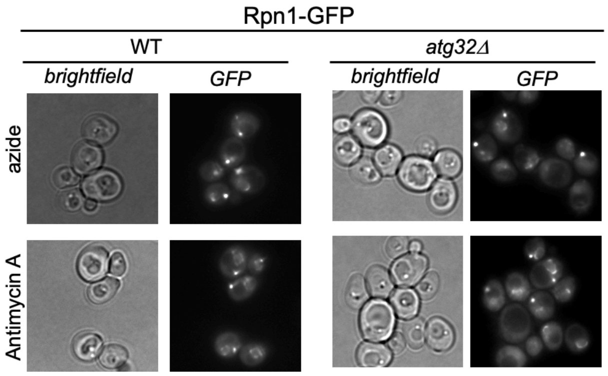


**S9.** Wild type (WT) and *atg32Δ* yeast expressing Rpn1-GFP were grown in rich raffinose media to log phase, treated with sodium azide or antimycin A and imaged after 24 hours. Data are representative of three independent experiments.

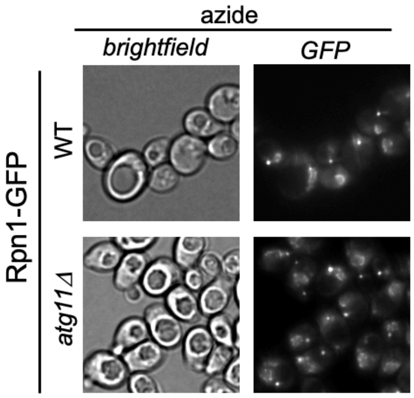


**S10.** Wild type and *atg11Δ* yeast expressing Rpn1-GFP were grown in rich raffinose media to log phase, treated with sodium azide and imaged after 24 hours. Data are representative of three independent experiments.

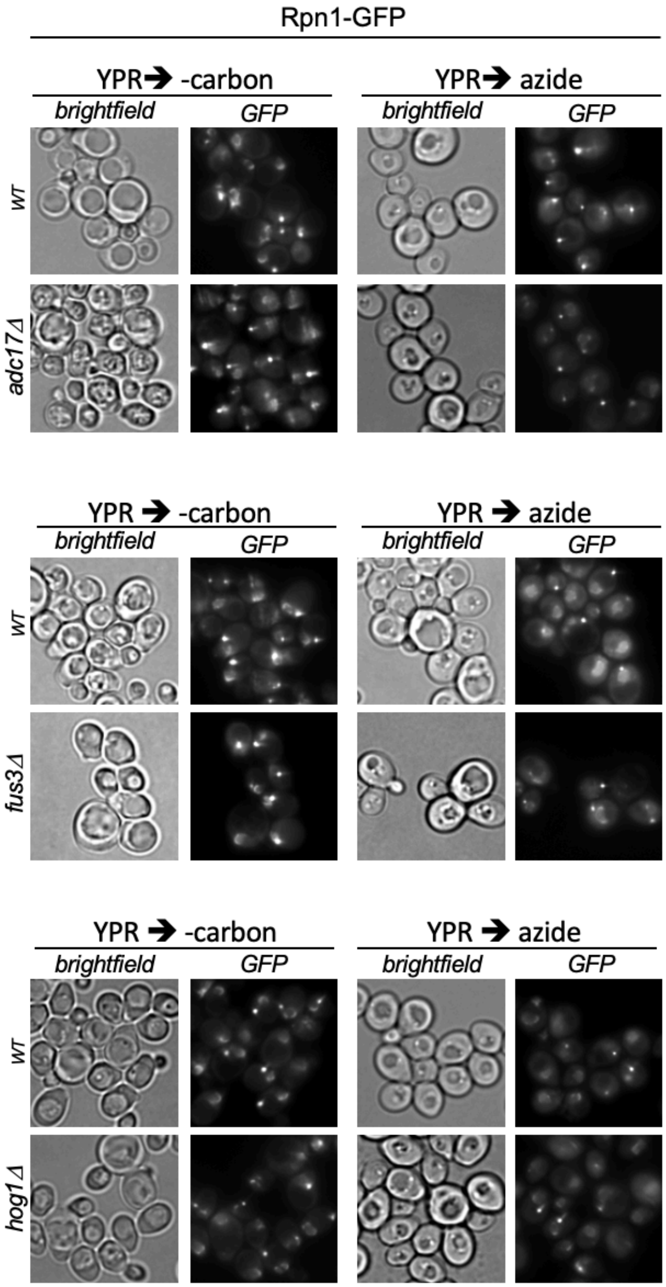


**S11.** Wild type, *adc17Δ* , *fus3Δ* and *hog1Δ* yeast expressing Rpn1-GFP were grown to log phase in rich media containing raffinose. Next, cells were starved for carbon or treated with sodium azide. Microscopy was performed at 24 hours. Data are representative of threeindependent experiments.

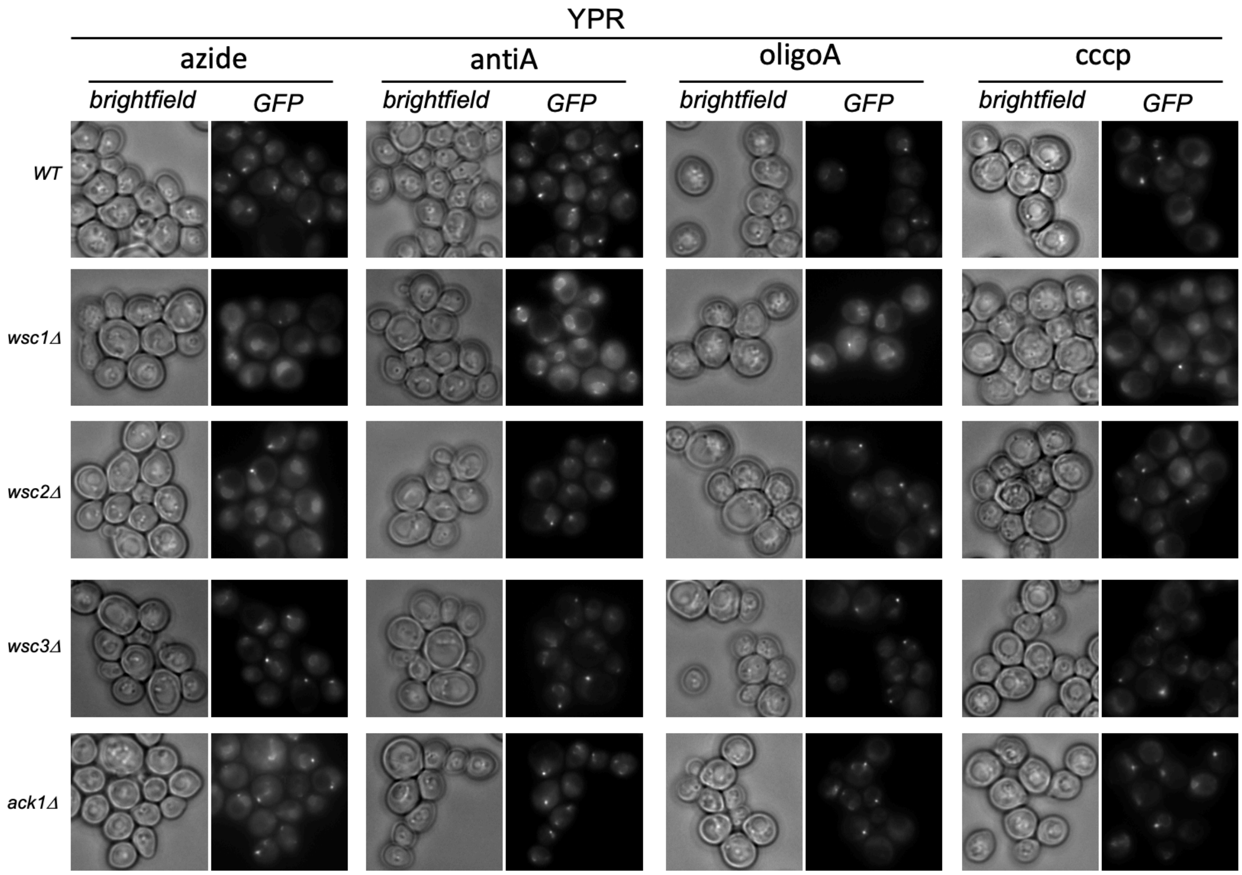


**S12.** Wild type, *wsc1Δ , wsc2Δ, wsc3Δ and ack1Δ* yeast expressing Rpn1-GFP were grown to log phase is rich media containing raffinose, and treated with mitochondria inhibitors for 24 hours before microscopy. Data are representative of two independent experiments.

**Blot Transparency:**

1A

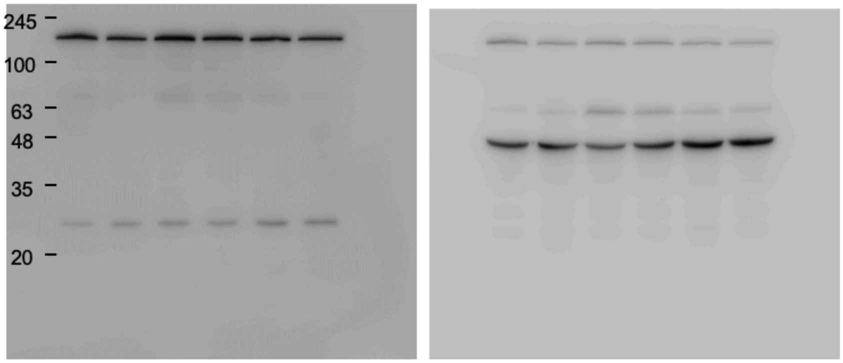


GFP Pgk1

1C/
S3

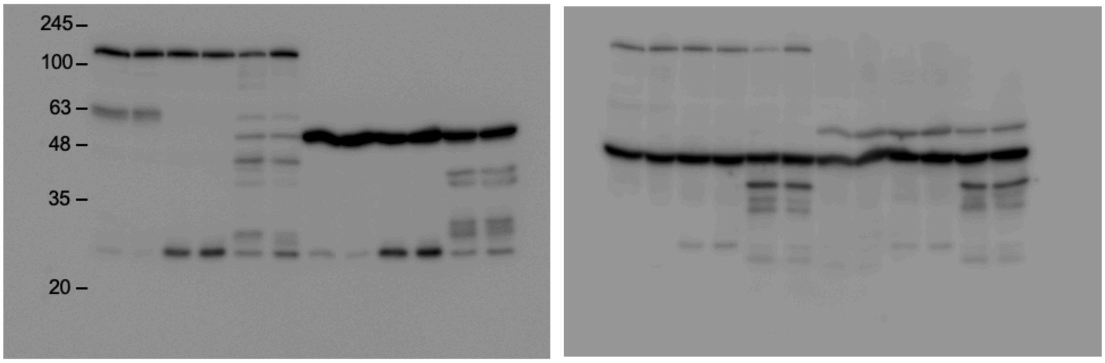


GFP Pgk1

4A

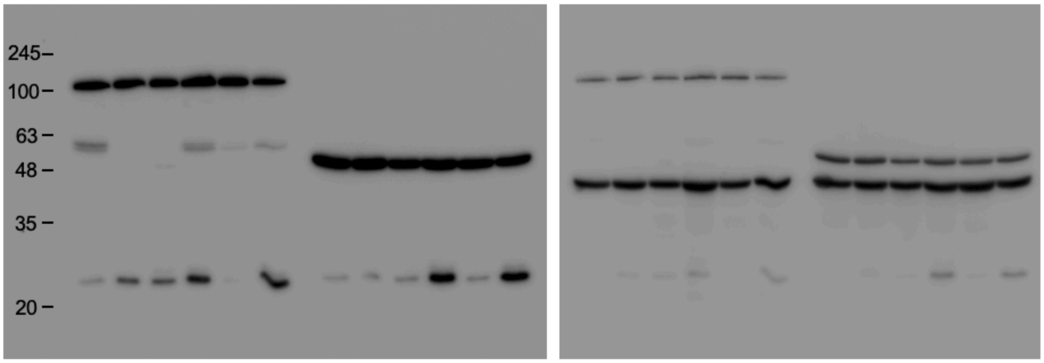


GFP Pgk1

**Supplementary table S1.** Strains used.

| Strain | Genotype (*lys2-801 leu2-3, 2-112 ura3-52 his3-Δ200 trp1-1)* | Figure | Source |
|---|---|---|---|
| SUB61 | MATα | **2e** | (1) |
| sJR861 | MATα *rpn1::RPN1-GFP (HIS3)* | **1a-c, 2a-d, 3a-g, 4a-c, S4, S6, S8-12** | (2) |
| sJR900 | MATα *rpn1::RPN1-GFP (HIS3) atg11:: G418* | **S10** | (2) |
| sJR1084 | MATα *scl1::SCL1-GFP (HIS3)* | **2c, 3a-g, 4a-c, S1-3, S7** | (3) |
| sJR1143 | MATA *rpn1::RPN1-mCherry (G418)* | **1d, e, S5** | (5) |
| sJR1151 | MATα *rpn1::RPN1-GFP (HIS3) mpk1::G418* | **3a-e** | (4) |
| sJR1216 | MATα *scl1::SCL1-GFP (HIS3) mpk1::HYG* | **3a-e** | (4) |
| sJR1224 | MATα *rpn1::RPN1-GFP (HIS3) fus3::G418* | **S11** | (5) |
| sJR1232 | MATα *rpn1::RPN1-GFP (HIS3) hog1::G418* | **S11** | (5) |
| sJR1264 | MATα *rpn1::RPN1-GFP (HIS3) ggc1::G418* | **S4** | (5) |
| sJR1270 | MATα *rpn1::RPN1-GFP (HIS3) snf1::G418* | **4a-c, S4** | (5) |
| sJR1271 | MATα *scl1::SCL1-GFP (HIS3) snf1::G418* | **4a-c** | (5) |
| sJR1308 | MATα *rpn1::RPN1-GFP (HIS3) adc17::G418* | **S11** | (4) |
| sJR1317 | MATA *ecm29::Ecm29-mCherry (G418) rpn1::RPN1-GFP (HIS3) atg1::HYG* | **S4** | (5) |
| sJR1327 | MATα *scl1::SCL1-GFP (HIS3) bck1:: G418* | **3g** | (4) |
| sJR1328 | MATα *rpn1::RPN1-GFP (HIS3) bck1:: G418* | **3g** | (4) |
| sJR1340 | MATα *rpn1::RPN1-GFP (HIS3) atg32:: G418* | **S4, S9,** | (5) |
| sJR1362 | MATα *rpn1::RPN1-GFP (HIS3) mig1:: G418* | **S3a** | (5) |
| sJR1392 | MATα *rpn1::RPN1-GFP (HIS3) mkk2::HYG mkk1::G418* | **3f** | (4) |
| sJR1393 | MATα *scl1::SCL1-GFP (HIS3) mkk2::HYG mkk1::G418* | **3f** | (4) |
| sJR1464 | MATα *rpn1::RPN1-GFP (HIS3) wsc1:: HYG* | **S12** | (5) |
| sJR1466 | MATα *rpn1::RPN1-GFP (HIS3) wsc2:: HYG* | **S12** | (5) |
| sJR1468 | MATα *rpn1::RPN1-GFP (HIS3) wsc3:: HYG* | **S12** | (5) |
| sJR1472 | MATα *rpn1::RPN1-GFP (HIS3) ack1:: HYG* | **S12** | (5) |

a) All strains have the DF5 background genotype ( lys2-801 leu2-3, 2-112 ura3-52 his3-**Δ**200 trp1-1)

1. Finley, D., Ozkaynak, E., and Varshavsky, A. (1987) Cell 48, 1035-1046
2. Waite, K.A., De La Mota-Peynado, A., Vontz, G., and Roelofs, J. (2015) JBC M115.699124
3. Waite, K.A., Burris, A, and Roelofs, J (2020) Sci. Rep. s41598-020-75126-1
4. Waite, K.A, Burris, A, Vontz, G, Lang, A, and Roelofs, J. (2021), JBC
5. This Study

**Supplementary table S2.** Primers used

| Primer | Genotype | Template | Sequence (5' to 3') |
| --- | --- | --- | --- |
| rvrs/rpn1 | *rpn1::rpn1-mCherry (G418)* | pBS34[3,] | TTTGAATTTTTCCTATTCTGGTTGATATTGCCCAAAAGCTATTCAGTTTAATCGATGAATTCGAGCTCG |
| pRL254 | *atg1::Hygro* | pFA6a-hph | TTC AAA TCT CTT TTA CAA CAC CAG ACG AGA AAT TAA GAA ACG TAC GCT GCA GGT CGA C |
| pRL265 | *atg1::Hygro* | pFA6a-hph | CAG GTC ATT TGT ACT TAA TAA GAA AAC CAT ATT ATG CAT CAA TCG ATG AAT TCG AGC TCG |
| pRL303 | *hog1::G418* | pFA6a-kanMX6 | CGGTAACCAGGCCATACAGTACGCTAATGAGTTCCAACAGCGTACGCTGCAGGTCGAC |
| pRL303 | *hog1::G418* | pFA6a-kanMX6 | ACATCAAAAAGAAGTAAGAATGAGTGGTTAGGGACATTAAATCGATGAATTCGAGCTCG |
| pRL401 | *rpn1::rpn1-mCherry (G418)* | pBS34[3,] | AGTAATTTTAAAGAAGAACCCTGACTATCGTGAAGAGGAGGGTCGACGGATCCCCGGG |
| pRL652 | *fus3::G418* | pFA6a-kanMX6 | CTACAAGGAAATAAGGCAGAGAAAAAGAAAGGAAAATAATCGTACGCTGCAGGTCGACG |
| pRL653 | *fus3::G418* | pFA6a-kanMX6 | ACATTGTTCTTCGGGTTGATATTTTAATGATAATGATGGCATCGATGAATTCGAGCTCG |
| pRL698 | *ggc1::G418* | pFA6a-kanMX6 | ATCAAACAAAGCATCTTCCAAAGTATTAGAAAAGGGAAATCGTACGCTGCAGGTCGACG |
| pRL699 | *ggc1::G418* | pFA6a-kanMX6 | AGCTGAATGCCAAGGAAAATAGACAAGAAGTTCATGTACTATCGATGAATTCGAGCTCG |
| pRL702 | *snf1::G418* | pFA6a-kanMX6 | TTTGTAACAAGTTTTGCTACACTCCCTTAATAAAGTCAACCGTACGCTGCAGGTCGACG |
| pRL703 | *snf1::G418* | pFA6a-kanMX6 | AAAAAAGGGAACTTCCATATCATTCTTTTACGTTCCACCAATCGATGAATTCGAGCTCG |
| pRL758 | *atg32::G418* | pFA6a-kanMX6 | ATCACAAAAGCAAAAAAAATCTGCCAGGAACAGTAAACAT CGTACGCTGCAGGTCGACG |
| pRL759 | *atg32::G418* | pFA6a-kanMX6 | GTGAGTAGGAACGTGTATGTTTGTGTATATTGGAAAAAGG ATCGATGAATTCGAGCTCG |
| pRL766 | *mig1::G418* | pFA6a-kanMX6 | CGAGAGTTGAGTATAGTGGAGACGACATACTACCATAGCC CGTACGCTGCAGGTCGACG |
| pRL767 | *mig1::G418* | pFA6a-kanMX6 | TCTTTTGATTTATCTGCACCGCCAAAAACTTGTCAGCGTA ATCGATGAATTCGAGCTCG |
| pRL853 | *wsc1::Hygro* | pFA6a-hph | GGCTGATTTAGTACTCAGGATAAAAAATTCTATTTAAATACGTACGCTGCAGGTCGACG |
| pRL854 | *wsc1::Hygro* | pFA6a-hph | GGCAATAGTTTAAGAATATAATAATTTTTTTTGGGTTTCTATCGATGAATTCGAGCTCG |
| pRL857 | *wsc2::Hygro* | pFA6a-hph | TTGAATTTCTTTTTGACCAACAGCATTATAGAAGTGGAATCGTACGCTGCAGGTCGACG |
| pRL858 | *wsc2::Hygro* | pFA6a-hph | GTCTTTGATATGAATATGTAGTGTTGGTATCTAAACCTAGATCGATGAATTCGAGCTCG |
| pRL861 | *wsc3::Hygro* | pFA6a-hph | TGTCCGTATAGTTTGTTTTTTTAGCAGAAGACAATATAAACGTACGCTGCAGGTCGACG |
| pRL862 | *wsc3::Hygro* | pFA6a-hph | CAGTTGATTACCATTAATGTTGAGAGTTTATGCGAGTTTTATCGATGAATTCGAGCTCG |
| pRL869 | *ack1::Hygro* | pFA6a-hph | TACCACCACGTTTTAATACTTTGTTTTAATACAGTAGGCACGTACGCTGCAGGTCGACG |
| pRL870 | *ack1::Hygro* | pFA6a-hph | TAATATTTGATTTATGGCATAATAAAGAAATGCGAATTCGATCGATGAATTCGAGCTCG |


1. Goldstein, A. L. & McCusker, J. H. Three new dominant drug resistance cassettes for gene disruption in Saccharomyces cerevisiae. Yeast 15, 1541–53 (1999).
2. Janke, C. et al. A versatile toolbox for PCR-based tagging of yeast genes: new fluorescent proteins, more markers and promoter substitution cassettes. Yeast 21, 947–62 (2004).
3. Hailey DW, Davis TN, Muller EG. Fluorescence resonance energy transfer using color variants of green fluorescent protein. Methods Enzymol. 351:34-49 (2002.
